# Narrow transmission bottlenecks and limited within-host viral diversity during a SARS-CoV-2 outbreak on a fishing boat

**DOI:** 10.1101/2022.02.09.479546

**Authors:** William W. Hannon, Pavitra Roychoudhury, Hong Xie, Lasata Shrestha, Amin Addetia, Keith R. Jerome, Alexander L. Greninger, Jesse D. Bloom

## Abstract

The long-term evolution of viruses is ultimately due to viral mutants that arise within infected individuals and transmit to other individuals. Here we use deep sequencing to investigate the transmission of viral genetic variation among individuals during a SARS-CoV-2 outbreak that infected the vast majority of crew members on a fishing boat. We deep-sequenced nasal swabs to characterize the within-host viral population of infected crew members, using experimental duplicates and strict computational filters to ensure accurate variant calling. We find that within-host viral diversity is low in infected crew members. The mutations that did fix in some crew members during the outbreak are not observed at detectable frequencies in any of the sampled crew members in which they are not fixed, suggesting viral evolution involves occasional fixation of low-frequency mutations during transmission rather than persistent maintenance of within-host viral diversity. Overall, our results show that strong transmission bottlenecks dominate viral evolution even during a superspreading event with a very high attack rate.

## Introduction

The long-term evolution of viruses is due to mutations that arise during replication within infected hosts and then transmit to new hosts. For viruses like SARS-CoV-2 or influenza that typically cause short self-limiting infections, evolution occurs over many consecutive rounds of infection, each interrupted by a transmission bottleneck. If there is a wide transmission bottleneck then mutations can gradually increase in frequency as a virus transmits from one host to another. However, a narrow transmission bottleneck means that low-frequency mutations present in a donor host will typically either be lost or fixed in a recipient host [1,2].

So far, efforts to understand how transmission shapes the evolution of SARS-CoV-2 have mainly focused on small household events or nosocomial pairs [3–7]. Such studies point to a narrow transmission bottleneck that significantly reduces viral genetic diversity at the start of each infection [3,4,6–8]. While exact estimates of the bottleneck range from 1 to 15 virions, it is clear that a limited number of virions initiate most human infections. These results are broadly similar to those for influenza, another heavily studied respiratory RNA virus [9–11].

However, it seems possible that the transmission of viral genetic diversity could show different patterns in different settings. For example, superspreading events play a significant role in SARS-CoV-2’s overall spread [12,13], and such events could exhibit different patterns of evolution since they involve settings highly conducive to viral transmission.

Here we investigate the spread of viral genetic diversity during a SARS-CoV-2 superspreading event on a fishing boat [14]. We perform high-depth metagenomic deep sequencing on nasal swabs collected from crew members of the fishing boat to characterize the intrahost populations of viral variants. Our results demonstrate that epidemiologically-linked individuals in a superspreading event share little to no intrahost viral diversity even at sites where mutations fix during the event, corroborating studies reporting narrow transmission bottlenecks in other settings [3,4,6–8].

## Results

### A large-scale SARS-CoV-2 transmission event on a fishing boat

We analyzed samples collected from an outbreak on a fishing boat in May 2020 [14]. There were a total of 122 individuals on the boat. Two days before embarking from Seattle, 120 individuals participated in pre-departure screening for SARS-CoV-2, and none tested positive. Despite this, infected crew members must have boarded the boat because a large SARS-CoV-2 outbreak ensued, eventually forcing the boat to return to shore in Seattle after 16 days at sea (Fig. 1A). Over 80% of crew members ultimately tested positive for SARS-CoV-2, indicating an extremely high secondary attack rate aboard the boat (Fig. 1B). Of note, only three crew members had neutralizing antibodies before the ship’s departure, and none of these individuals met the case definition for infection [14].

**Figure 1.**
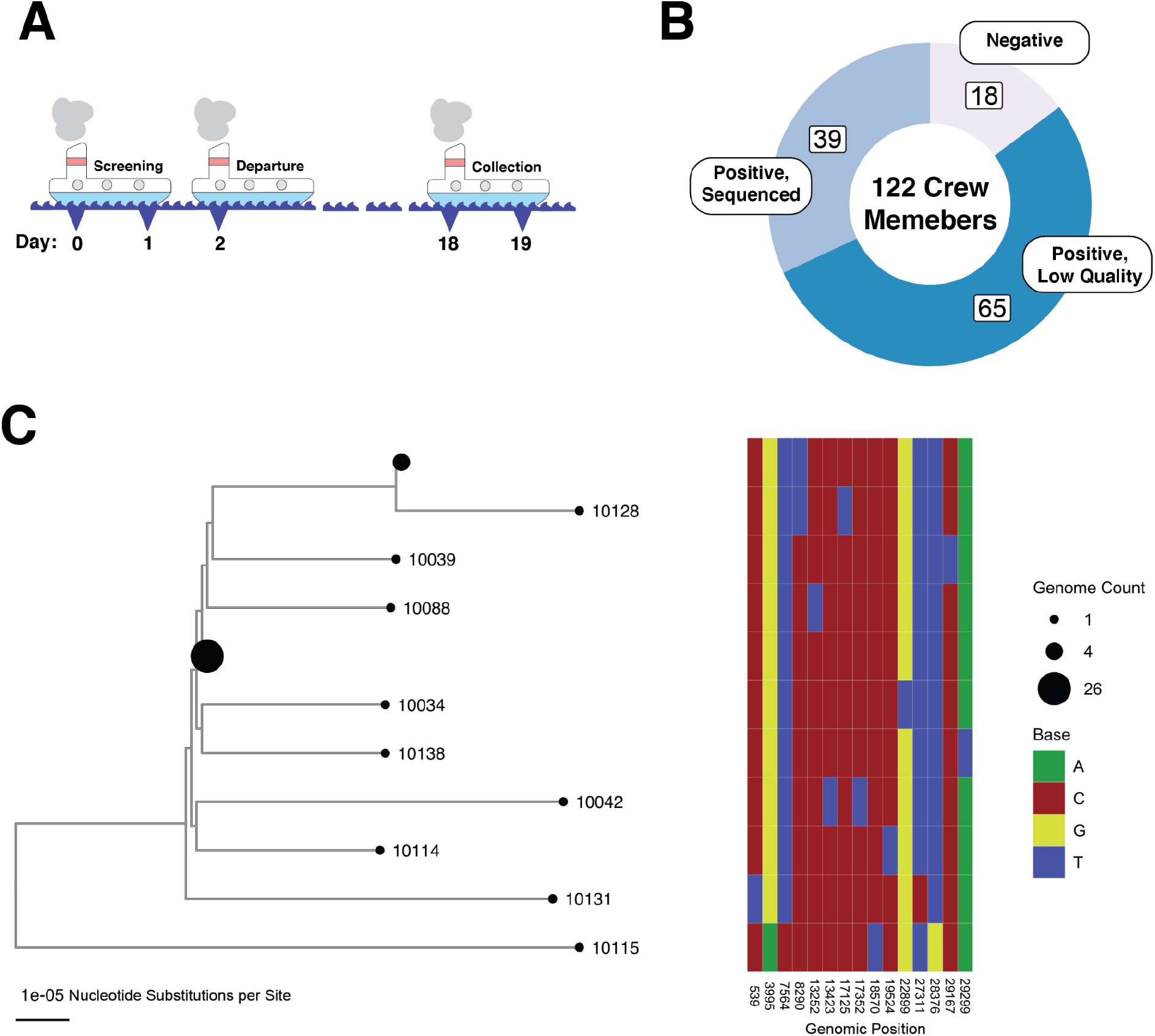
An outbreak of SARS-CoV-2 on an isolated fishing boat is an epidemiologically linked cluster of infections. (A) Schematic showing the timeline of the fishing vessel outbreak. All samples used in this study were taken on day 18 as shown in the figure (relative to the start of pre-departure screening). (B) Donut plot showing the sampling breakdown for all 122 members of the crew. (C) Phylogeny of SARS-CoV-2 genome from the boat. A heatmap to the right shows the nucleotide differences between genomes on the tree. Specimen identification numbers for crew member samples label the leaf nodes of the tree except for those nodes with more than one identical genome. Node sizes are proportional to number of sequences: there is a node representing 26 identical sequences(10101, 10126, 10133, 10105, 10108, 10130, 10031, 10110, 10030, 10124, 10029, 10102, 10038, 10094, 10027, 10118, 10117, 10106, 10091, 10093, 10127, 10116, 10040, 10090, 10036, 10089)and a node representing 4 identical sequences (10107, 10129, 10113, 10028); all other nodes represent unique sequences.

Nasal swabs were collected from the crew members two days after the boat returned to shore. Of the samples that were positive in a SARS-CoV-2 PCR test, 39 had sufficiently high levels of viral RNA (Ct value less than 26) to assemble consensus viral sequences from deep sequencing data, as previously described in Addetia et al (Fig. 1B). These consensus viral sequences from the boat samples differed on average at fewer than two positions, and were clearly diverged relative to the non-boat outgroup sample (Fig 1C). Over 75% of the viral sequences from the boat were identical to at least one other sequence from the boat. Given the genetic similarity of viral sequences from the boat and the short time frame for infections, this cohort resembles a superspreading event.

To place the superspreading event in the larger context of SARS-CoV-2’s genetic diversity, we inferred a phylogeny using representative sequences from viruses circulating before the outbreak, including a subset of the most genetically similar viral sequences to those isolated from the boat. The boat clade is nearly monophyletic, although two surveillance sequences collected elsewhere in Washington state around the time of the outbreak fall in the same clade as the boat samples (Fig. 2). These sequences likely share a close common ancestor with the virus that seeded the superspreading event on the boat. We also chose one Washington state sample not from the boat for further sequencing, and as expected this sample was distinct from the boat clade on the tree. Overall, the nearly monophyletic nature of the outbreak clade and the fishing boat’s isolation makes this cohort appropriate for assessing how SARS-CoV-2 genetic diversity transmits among a tightly associated group of individuals.

**Figure 2.**
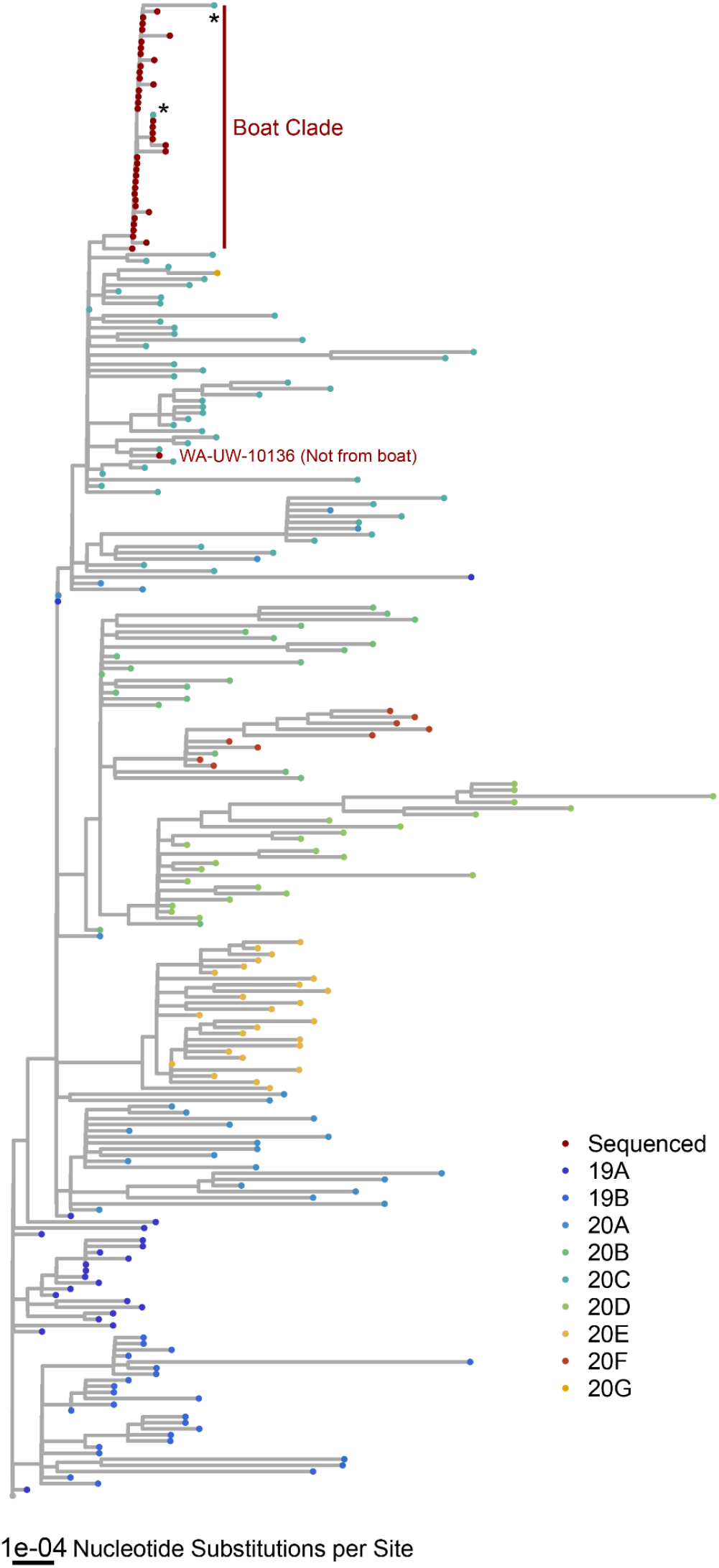
Sequences from the boat form a distinct clade. A phylogeny of the 39 crew-member genomes and representative genomes from other circulating clades before the outbreak. Additionally, this phylogeny includes the ten closest matches to each of the 39 crew-member genomes from a custom BLASTN database made with sequences collected from Washington in a two-month interval around the time of the outbreak. We also re-sequenced as a control one sample not from the boat (WA-UW-10136). Most genomes isolated from the boat form a distinct clade broken only by two genomes (hCoV-19/USA/WA-UW-10510/2020 and hCoV-19/USA/WA-UW-10521/2020) annotated with an asterisk.

### High-quality deep sequencing of samples with adequate viral RNA

We used deep sequencing to measure the intra-host viral genetic variation in the samples collected from infected crew members. We employed several approaches to ensure the accuracy of these measurements. First, we used a shotgun metagenomic sequencing approach to avoid potential mutational biases from specific PCR amplification of viral RNA. Of the 39 nasal swabs described in the previous section, 23 had sufficient viral RNA (Ct value less than 20) to sequence metagenomically (Supplemental Table 1) [15]. Second, we sequenced replicates starting from independent reverse-transcription reactions from the same initial nasal swab. In principle, each replicate should sample from the same underlying viral population, and so differences between replicates can indicate limitations due to a lack of underlying template molecules in the swabs [16,17]. We used a stringent cutoff for sequencing depth by only considering sequences with >80% of the genome covered by 100 reads in one or more replicates in the downstream analysis (Fig. S1A). There were no biases observed in sequencing coverage across the length of the viral genome (Fig. S2A).

We compared results between replicates for each crew member and focused our subsequent analyses on the 13 crew members with high concordance between replicates and adequate sequencing depth (Fig. 3). Of note, the results were robust to using different methods for variant calling (Fig S2 and S3).

**Figure 3.**
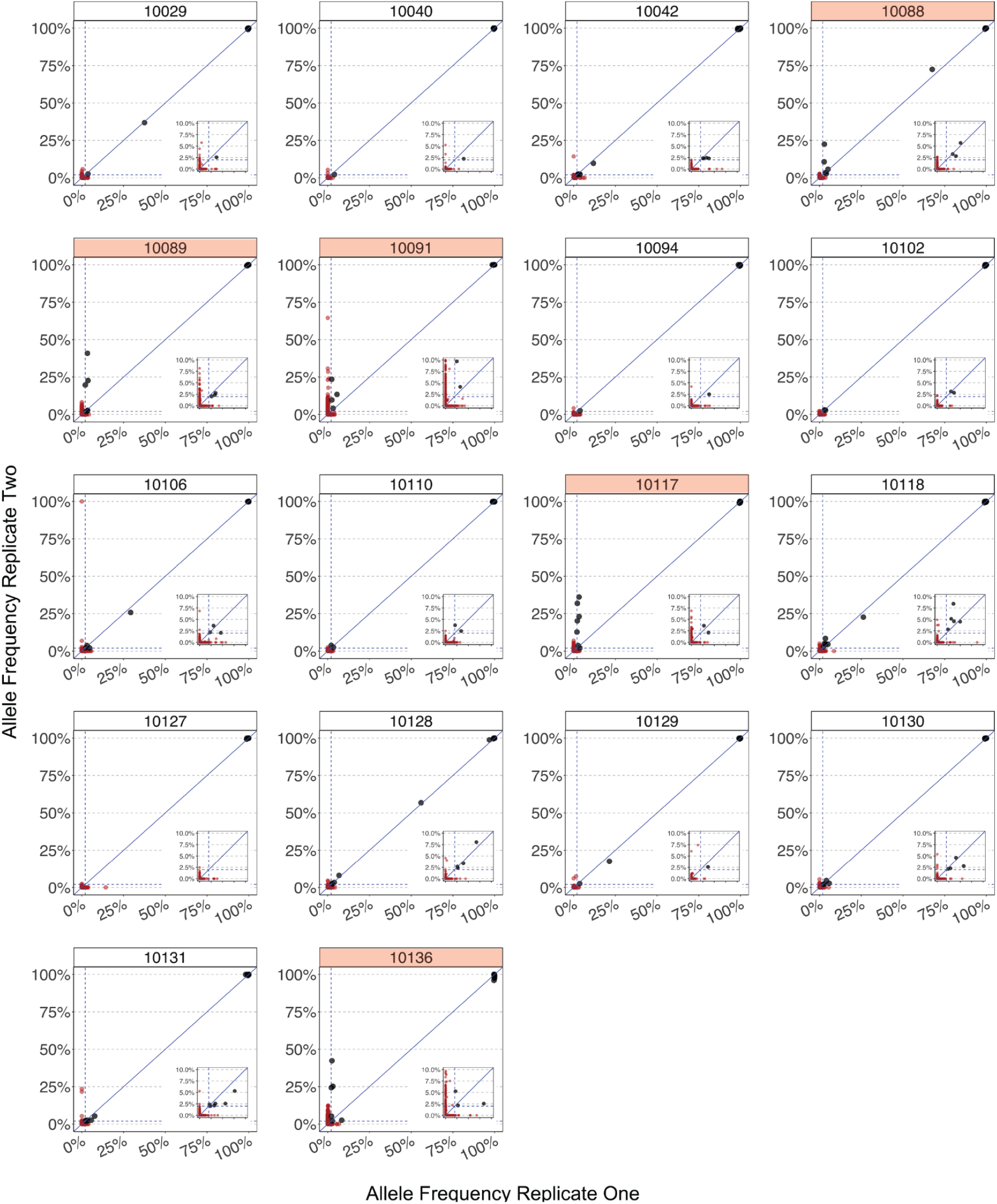
Robust quality control reveals false-positive variant alleles and samples of poor quality. Each plot shows the concordance between allele frequencies between replicates for every specimen that we sequenced, with both replicates having greater than 100X coverage in at least 80% of the genome. Alleles that were present in less than 2% of 100 reads in either replicate are colored red. The dotted line represents the 2% frequency threshold. We highlighted the facet headers of ‘poor’ quality crew member samples in red if there was a large discrepancy in allele frequencies between replicates. This figure also shows the non-boat sample (10136) sequenced as a control.

### The intrahost virus population is relatively homogeneous

After retaining just the samples with high sequencing depth and good replicate-to-replicate correlations, we assembled a set of intrahost SNPs that were present in 2% of at least 100 reads in both replicates. To determine the extent of within-host diversity in each patient, we converted any mutation (relative to the reference) above 50% frequency to its corresponding minor allele and counted the total number of minor allele variants at >2% frequency per crew member. The diversity of the virus populations within each crew member was limited, with an average of three intra-host variants per individual (range 0-5, Fig. 4A). Furthermore, most intra-host variants were at relatively low frequencies, with only a handful at >10% (Fig. 4B). This limited within-host diversity and low-frequency-dominated allele frequency spectrum are consistent with other studies of SARS-CoV-2 intrahost diversity that have utilized robust computational and experimental controls (Fig. 4B) [3,4,8,18]. There was no correlation between the Ct value of the nasal swab and the number of SNPs we identified (Fig. S3). Additionally, there was no discernable pattern in the location of SNPs in the genome (Fig S4 and S5).

**Figure 4.**
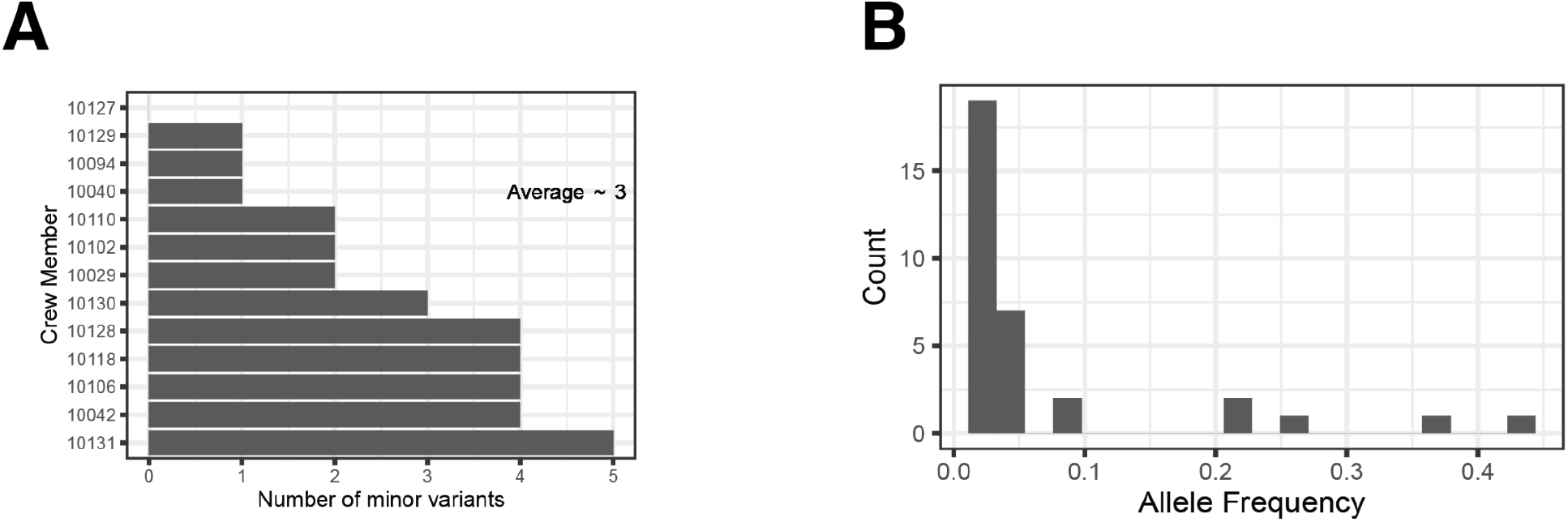
The intra-host spectrum of minor alleles reveals a relatively homogeneous virus population. (A) Bar graph showing the number of minor variants (< 50% allele frequency) identified in both replicates of each crew member. There was an average of three minor variants per infection across the ten crew members. (B) The minor allele frequency spectrum across all twelve crew member specimens with minor variants.

### Mutations that fix on the boat are not observed at intermediate frequencies

We next considered two possible models for how mutations could spread and fix on the boat. The first model assumes that the transmission bottleneck is narrow, and variants will either be lost during transmission or, less frequently, they will fix during a single transmission event. The second model assumes that the transmission bottleneck is wide, and variants will transmit between multiple infections and gradually rise in frequency until they fix (Fig. 5A).

**Figure 5.**
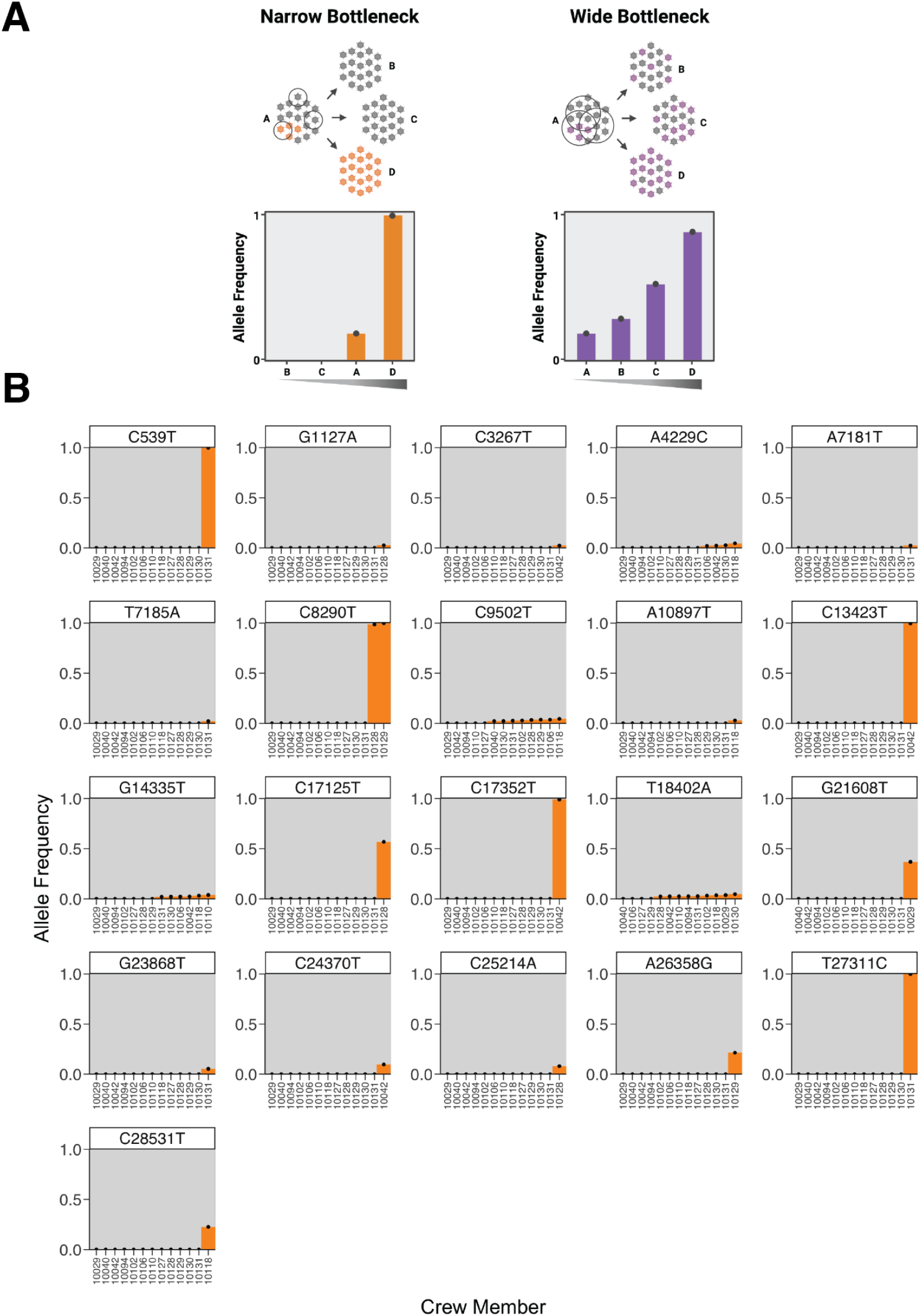
The spectrum of shared minor variation suggests that the transmission bottleneck is narrow. (A) A schematic showing the expected pattern of observed allele frequencies for shared variants in either a narrow or wide bottleneck scenario. (B) Each plot represents the frequency of a single nucleotide polymorphism (SNP) across crew members. Variants are called relative to the ancestral sequence of the virus introduced to the boat as inferred from the phylogeny of crew member genomes. The x-axis is ordered by variant frequency.

To determine which model best describes viral transmission on the boat, we plotted the frequency of every variant allele for each crew member and sorted the crew members by allele frequency. We identified variants relative to the inferred ancestral sequence for the root of the boat clade (which is also the consensus and most common sequence on the boat, see Fig. 1A). If the transmission bottleneck is narrow, most non-fixed variants would be private to single individuals, and at sites with fixed variants the mutations will generally be present at ~0% or ~100% frequency. If transmission bottlenecks were wide on the boat, variants would be observed in multiple individuals at intermediate frequencies. We observed that most low-frequency variants were private to single individuals, and fixed variants were never also observed at intermediate frequencies (Fig. 5B). The lack of a gradient in the frequency for fixed variants on the boat suggests that viral evolution on the boat is dominated by a narrow transmission bottleneck.

Although most variants were either fixed or private to single crew members, four low-frequency alleles were present in multiple individuals on the boat (A4229C, C9502T, G14335T, and T18402A in Fig. 5B). However, none of these variants ever reached more than 5% frequency. Furthermore, several characteristics of these shared low-frequency variants suggest they are sequencing artifacts rather than true mutations. First, these same variants are also observed in our deep sequencing of a control sample not collected from the boat but sequenced in the same run as the boat samples (Fig. S6). Furthermore, one variant, C9502T, is present in a homopolymeric stretch of thymines, a known correlate with spurious variant calls in SARS-CoV-2 sequencing data [3,19]. Additionally, G14335T and A4229C exhibit significant positional bias in the aligned reads, with most observations at the beginning of the read. Read position correlates with false-positive variant calls in experimental studies of viral deep-sequencing data [20]. Finally, T18402A demonstrates significant divergence in its frequency between replicates. These four shared variant alleles are therefore likely technical artifacts that survived our quality checks.

## Discussion

This study examined the spread of SARS-CoV-2 genetic diversity during a superspreading event on a boat. We found low rates of intrahost viral diversity among infected individuals, and mutations that did fix appeared to do so during single transmission events. Our results demonstrate that transmission of intrahost viral diversity is limited even during superspreading events that are highly conducive to transmission. These findings are consistent with studies of SARS-CoV-2 transmission in other settings such as households or hospitals [3,4,6–8], suggesting narrow transmission bottlenecks are a near universal feature of the virus’s transmission. Similar narrow transmission bottlenecks also dominate the evolution of influenza virus [9–11].

A key aspect of our study was sequencing duplicates and rigorous variant calling. False-positive variants shared between multiple samples significantly biased the results of Popa et al., leading to an estimate of the bottleneck nearly 10-fold higher than other studies [5,8]. Martin and Koelle reanalyzed this data with a more stringent allele frequency filter, and the bottleneck estimate dropped from greater than 1000 founding viruses to between 1 and 3 founding viruses [8]. Despite our attempts to remove low-frequency false-positive variants, some survived our quality controls. Further research to determine the cause of shared false-positive variants in clinical SARS-CoV-2 deep sequencing could further improve the accuracy of these studies.

Our study has several limitations. First, we were able to obtain high-quality sequencing for only some of the boat’s crew members. After accounting for samples that passed our quality controls, only 13 of the 122 crew members were available for analysis. Therefore, we might be missing instances where a variant rises to fixation over multiple transmission events. Another limitation of this study is that we cannot quantitatively estimate the transmission bottleneck because we do not know which passengers infected one another. However, we can still observe that the paucity of shared intra-host viral variants indicates generally narrow transmission bottlenecks.

Overall, our study corroborates the finding of limited shared intra-host viral diversity that has been observed in studies of acute infections with SARS-CoV-2 in other settings. Therefore, even superspreading events in poorly ventilated, close-quarters environments appear insufficient to alter the dominant role of transmission bottlenecks in shaping the evolution of SARS-CoV-2.

## Methods

### Ethics Statement

The use of residual clinical specimens was approved by the University of Washington IRB (protocol STUDY00000408) with a waiver of informed consent.

### Sample Collection and Preparation

RNA was extracted from positive SARS-CoV-2 nasal swabs from crew members using the Roche MagNa Pure 96 [21]. The initial sequencing libraries were constructed as previously described and sequenced on a 1 x 75 bp Illumina NextSeq run [14]. RNA was DNase treated using the Turbo DNA-Free kit (Thermo Fisher). First-strand cDNA synthesis was performed using Superscript IV (Thermo Fisher) and 2.5 μM random hexamers (Integrated DNA Technologies), and second-strand synthesis with Sequenase version 2.0 DNA polymerase (Thermo Fisher). Double-stranded cDNA was purified using AMPure XP beads (Beckman Coulter) and libraries were constructed using Nextera Flex DNA pre-enrichment kit with 12 cycles of PCR amplification (Illumina). We re-sequenced samples from these original libraries to increase their depth if they had a RT-qPCR Ct values less than 20 from an RT-qPCR as measured in a previous paper [14]. Samples with a Ct value less than 20 were deemed to have enough RNA to be sequenced without specific amplification of viral RNA by PCR with targeted primers.

Additionally, we made duplicate libraries starting from the same nasal swabs as the initial library using independent reverse transcription reactions and identical library preparation methodology. In principle, each replicate should sample from the same underlying virus population, so differences between replicates can indicate limitations due to a lack of underlying template molecules in the swabs [11,17]. Of note, one specimen from the original paper, 10136, was subsequently determined not to have been isolated from the boat but general viral surveillance in Seattle. We kept this sample and resequenced it as non-boat control. We obtained an average of 1,113,690 mapped reads per library.

### Sequencing Data Processing

All data processing from the raw unaligned sequencing files onwards was handled by our Snakemake pipeline available on Github — https://github.com/jbloomlab/SARS-CoV-2_bottleneck [22]. Sequencing reads from the raw FASTQ files from each sequencing run were trimmed for adaptor sequences and long (>10) homopolymer sequences at the ends of reads with fastp [23]. Fastp was also used to filter reads from the FASTQ file if they contained more than 50% unqualified bases (Phred < 15) or were less than 50 base pairs in length. Following quality filtering, SARS-CoV-2 specific reads were selected from the FASTQ files by matching 31 base long kmers to the Wuhan-1/2019 reference genome (NC_045512.2) using BBDuk (https://jgi.doe.gov/data-and-tools/bbtools/bb-tools-user-guide/bbduk-guide/).

After quality filtering and selection of reads containing SARS-CoV-2 sequences, the FASTQ files were aligned to the Wuhan-1/2019 reference (NC_045512.2) with BWA mem [24]. Libraries that were resequenced for greater depth were joined these libraries together after alignment with Samtools merge [24]. The aligned BAM files were checked for quality using Samtools to obtain average coverage, base quality, and completeness.

### Phylogenetic Analysis

We used aligned BAM files to make consensus sequences for each crew member. Individual consensus sequences were created for each replicate by taking the most represented base at every position if that position had more than 100 reads with a base quality score of greater than 25, otherwise, an N was added to the sequence. Then, we combined the consensus sequences from each replicate and filled in Ns where possible. If the consensus from each replicate disagreed at a position, an N was inserted. In addition to the consensus genomes from 23 crew member samples we deep-sequenced in duplicate, we included 16 consensus genomes from the boat assembled in the previous study downloaded from GISAID [14]. Following the assembly of consensus genomes for each crew member, we aligned the genomes with MAFFT [25]. We masked the non-coding 3’ and 5’ portions of the genome. Using these aligned genomes, we built a phylogenetic tree with IQtree using 1000 bootstrap iterations with an invariable site plus discrete gamma model and ancestral state reconstruction [26,27]. The ancestral state reconstruction was used to infer the ancestral sequence of the genomes from the boat. The tree was rooted using midpoint rooting as implemented by the R package phytools and plotted with ggtree [28,29] (Fig. 1C).

To determine where all of the available boat sequences fit in the coincident global phylogeny, we downloaded at most 25 genomes from GISAID that met our quality criteria (<5% Ns, high coverage, complete coverage, and human host) from each of the circulating Nextrain clades at and before the time of the outbreak on May 5th, 2020 (19A, 19B, 20A, 20B, 20C, 20D, 20E, 20F) [30]. Additionally, to include genomes that were similar to those on the boat, we built a BLASTN database from all sequences collected in Washington state at and before the time of the outbreak (May 5th, 2020) that met the same quality standards described above. We took the 10 closest matches to each of the 24 consensus genomes to include in the phylogeny. We aligned these sequences using MAFFT, however, we also aligned to the Wuhan-1/2019 (NC_045512.2) reference and standardized the length of each sequence. Following alignment, we masked the sequence before the start of ORF1ab and after position 29675 to control for sequencing errors at the start and end of the genome. We built a phylogeny with IQtree using the same parameters as above. The tree was rooted using outgroup rooting with the Wuhan-1/2019 reference (NC_045512.2) as the outgroup as implemented by the R package ape and plotted with ggtree [28,31] (Fig. 2).

The code to run all of the phylogenetic analyses is provided on Github at https://github.com/jbloomlab/SARS-CoV-2_bottleneck/blob/master/workflow/notebooks/Phylogenetic-Analysis.ipynb. The GISAID IDs for sequences used to conduct this analysis are listed in the supplement along with their submitting lab (Supplemental Table 2).

### Variant Calling and Filtering

Variants were identified using a custom Python script (https://github.com/jbloomlab/SARS-CoV-2_bottleneck/blob/master/workflow/scripts/process_pysam_pil_eup.py). Briefly, we counted the coverage of each base at every position in the reference genome using the python/samtools interface Pysam (https://github.com/pysam-developers/pysam). Bases were only included if they surpassed a Phred quality score of 25. After identifying SNPs, our program goes back through the BAM file and identifies reads that overlap these polymorphic sites. We record the total number of occurrences of the SNP, the average position in each read, and the strand ratio. SNPs present after position 29860 in the genome were excluded from the output to avoid sequencing artifacts. The final SNPs were annotated for coding effect and position in the genome using another custom script (https://github.com/jbloomlab/SARS-CoV-2_bottleneck/blob/master/workflow/scripts/annotate_coding_changes.py).

In addition to our custom approach, we also called variants using three different variant calling programs, ivar, varscan2, and lofreq [32–34]. Where applicable, the same heuristic filters were used in each program. The minimum base quality score was 25, the minimum coverage was 100X, at least 10 reads needed to contain a given SNP, and the minimum allele frequency was 0.5%. Filters that could not be applied in a given program were standardized post-hoc in R. Variants from each program were standardized into a similar format and added to a single table. Insertions and deletions were removed as we did not benchmark our pipeline to detect these. We annotated the coding effect of each SNP using SnpEff [35]. These extra sets of shared variants were used to cross-check the results of our approach with that of others.

Finally, to identify variants that were shared between individuals on the boat and determine how variants came to be fixed (Fig. 5), we considered all variants relative to the ancestral boat sequence inferred by IQtree using a phylogeny of the boat sequences. Therefore, the included fixed mutations arose after the introduction of the virus to the boat.

## Data Availability

All sequencing data are available on the NCBI SRA at the project accession PRJNA803551. All code used to run the analyses described in this paper are archived on Github (https://github.com/jbloomlab/SARS-CoV-2_bottleneck) and Zenodo at *doi.xxxxx.*

## Acknowledgements

The work in the lab of JDB was supported in part by NIH / NIAID (R01AI141707). JDB is an Investigator of the Howard Hughes Medical Institute. This project has been funded in part with Federal funds from the National Institute of Allergy and Infectious Diseases, National Institutes of Health, Department of Health and Human Services, under Contract No. 75N93021C00015. PR is a CFAR New Investigator award recipient supported by NIH AI027757. We acknowledge all authors from originating and submitting laboratories of the sequences from GISAID’s EpiCoV database. An acknowledgments table is available in the Supplementary Materials.

## Supplement

**Supplementary Figure 1.**
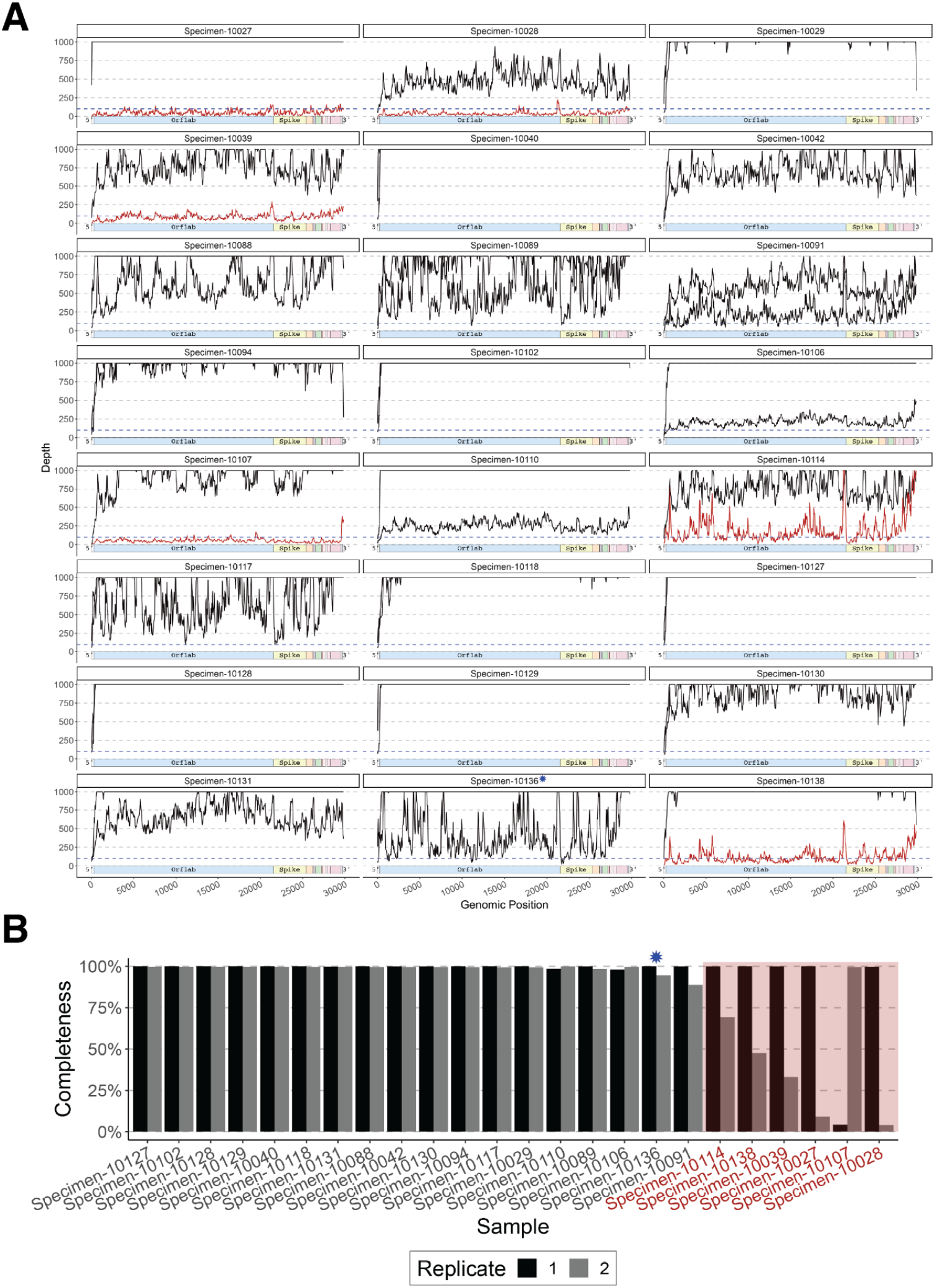
Samples were filtered by completeness. (A) The pattern of sequencing depth for each replicate of all 23 specimens from the boat, and one control sample that was not from the boat (Specimen-10136, labeled with a blue asterisk), that we resequenced for this study. The number of reads per site is capped at 1000X coverage. Samples colored in red have less than 80% of the genome covered by 100X reads. (B) Completeness refers to the percentage of the genome covered by more than 100X reads. Samples colored and highlighted in red have at least one replicate with less than 80% of the genome covered by 100X reads. These samples were excluded from the downstream variant analysis.

**Supplementary Figure 2.**
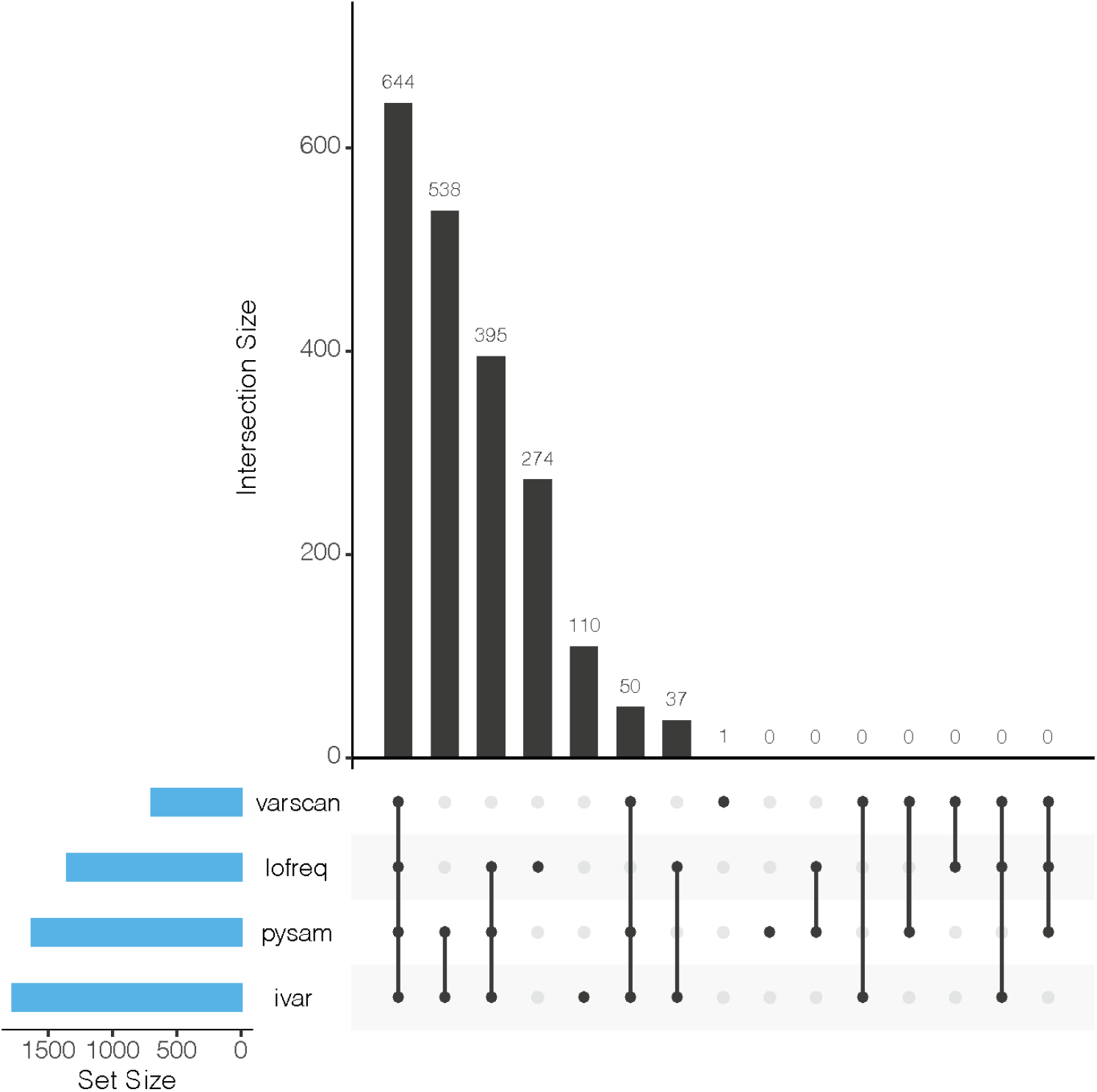
Comparison between different variant calling methods. An UpSet plot shows the overlap in the sets of SNPs called by three different variant calling methods - varscan2, lofreq, ivar, and our custom python script using pysam *(Citations).* Variants were covered by more than 100X reads and present at greater than 2% frequency to be included in the set for each variant caller. The majority of variants are called by all four methods. No variants are called by our custom script that aren’t identified by at least one other method.

**Supplementary Figure 3.**
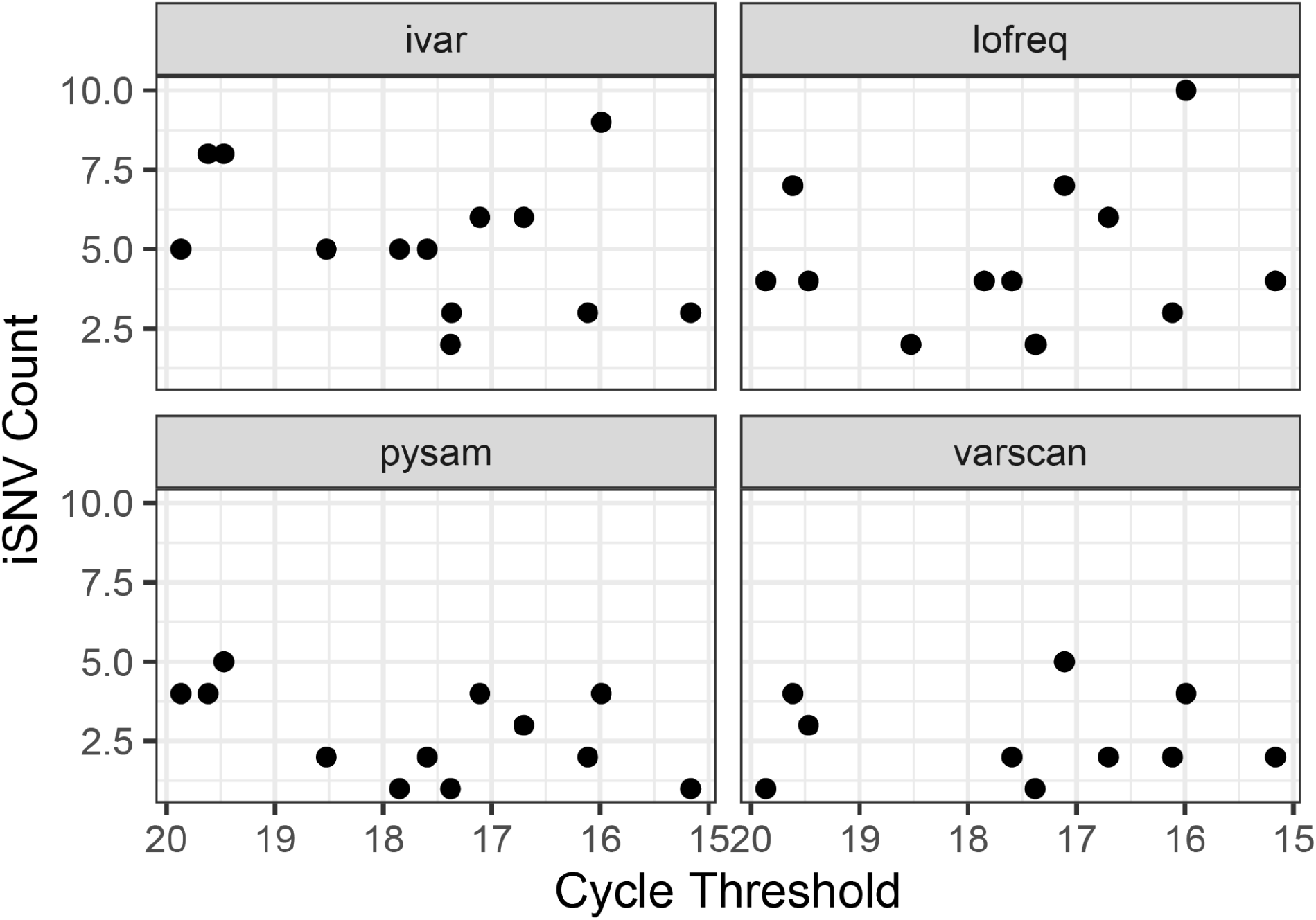
Ct value does not correlate with the number of polymorphisms. Regardless of the variant calling method used, the Ct value of the original nasal swab does not correlate with the number of variants called after filtering out low-frequency (>2%) and poorly covered (>100X) variants. Only samples that passed our quality controls for sequencing completeness (Fig. S3B) and concordance (Fig. 2) were included in this analysis.

**Supplementary Figure 4.**
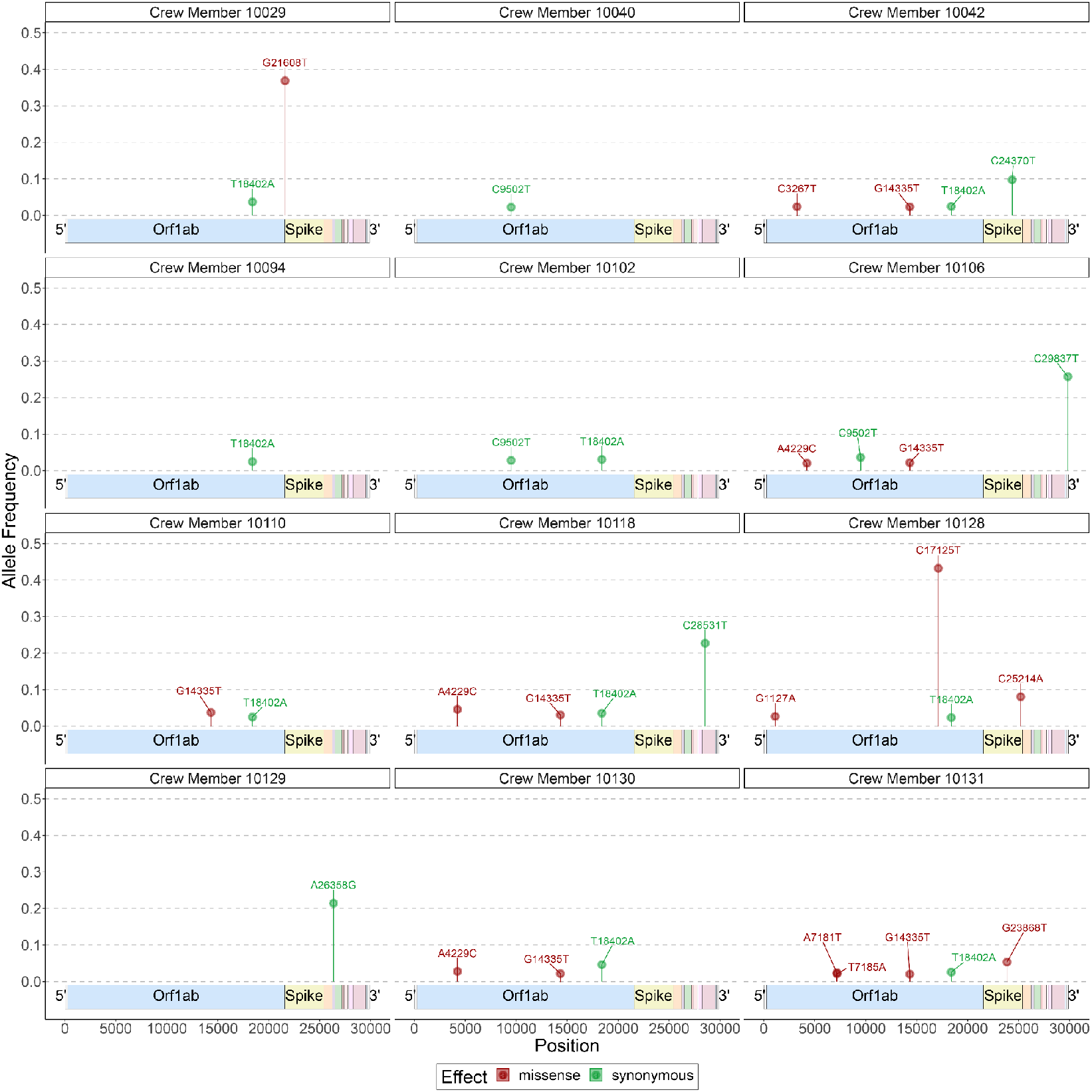
There is no discernable pattern of minor variants in the genome. Plot showing every minor variant (>50% allele frequency) identified across the crew members that passed our quality filters. We included variants if they were present in more than 2% of greater than 100 reads.

**Supplementary Figure 5.**
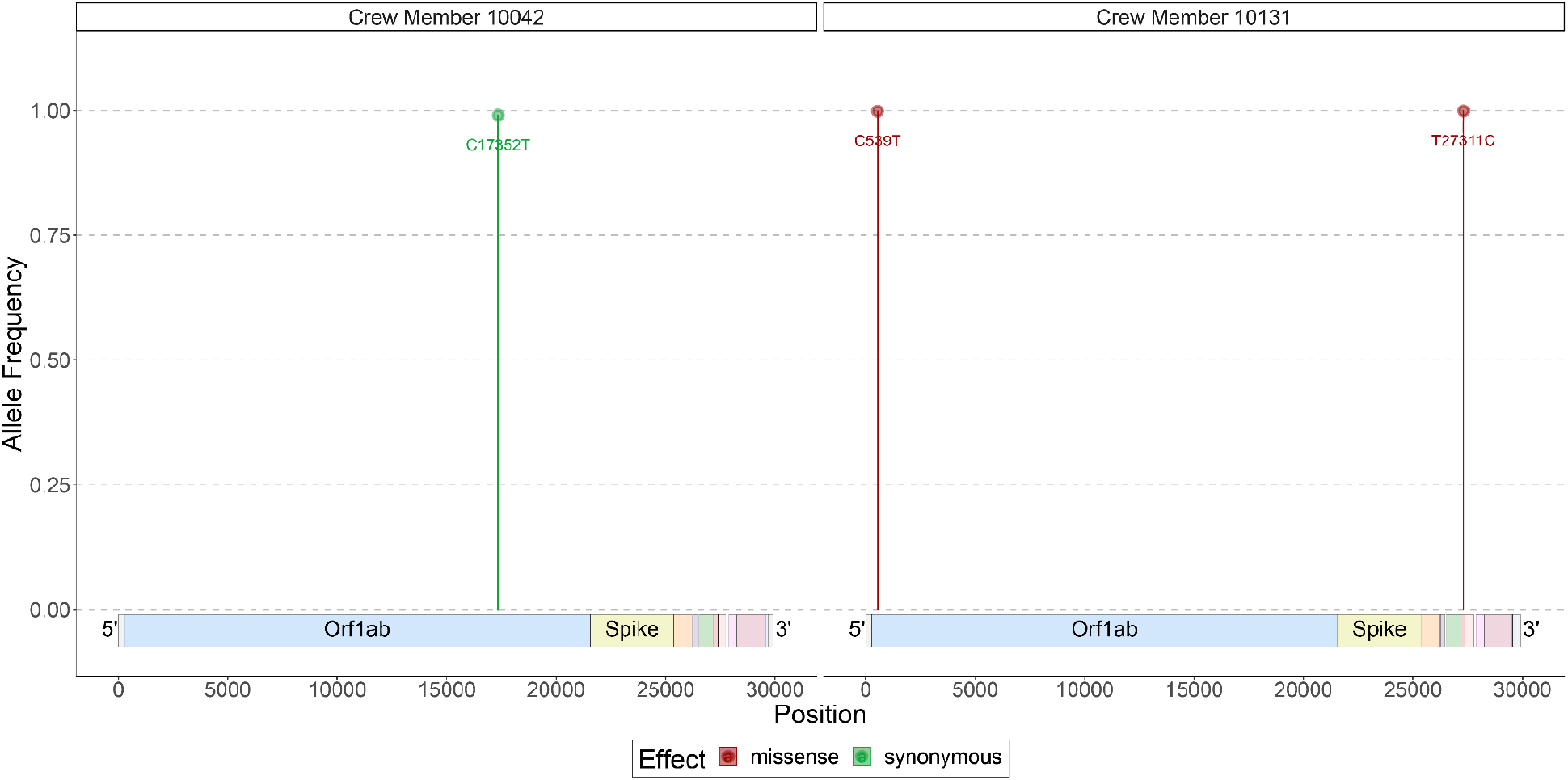
Distribution of fixed mutations in the genome. Plot showing fixed variants identified across the crew members that passed our quality controls. We included variants if they were present in 98% or more of at least 100 reads. Mutations that are present in the 5’ and 3’ UTRs are excluded from this plot.

**Supplementary Figure 6.**
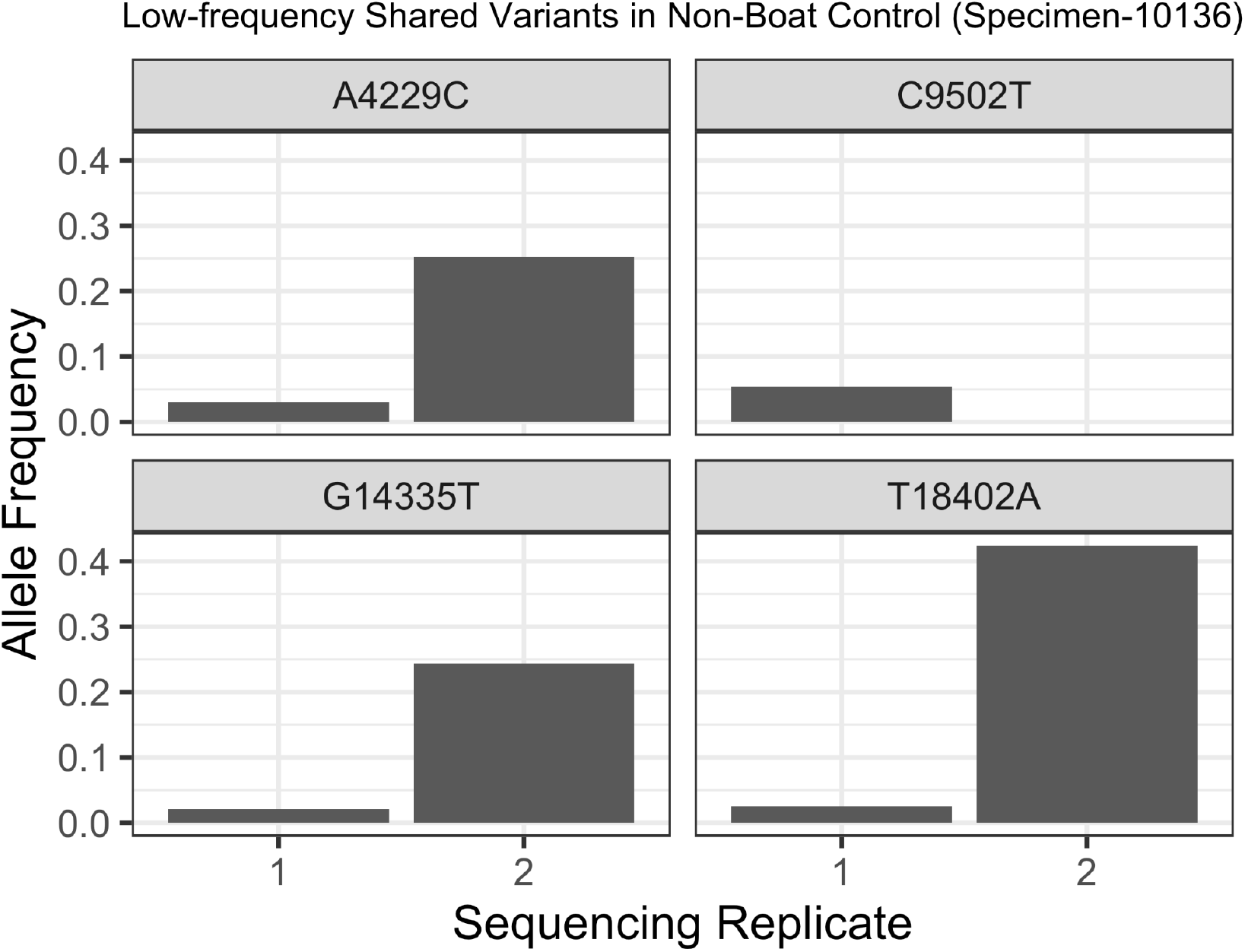
Low frequency shared variants are present in the non-boat control specimen. Four variants shared at low-frequency between crew members are also detected in a specimen not collected from the boat but included as a control in both sequencing runs (Specimen 10136). This observation suggests that these are not de novo low-frequency variants that arise on the boat and spread between the crew, but rather sequencing contamination or variant calling errors common to samples from the two sequencing runs.

